# Long-Distance Electrical and Calcium Signals Evoked by Hydrogen Peroxide in Physcomitrella

**DOI:** 10.1101/2023.04.21.537805

**Authors:** Mateusz Koselski, Sebastian N. W. Hoernstein, Piotr Wasko, Ralf Reski, Kazimierz Trebacz

## Abstract

Electrical and calcium signals in plants are one of the basic carriers of information transmitted over a long distance. Together with reactive oxygen species (ROS) waves, electrical and calcium signals can participate in cell-to-cell signaling, conveying information about different stimuli, e.g. abiotic stress, pathogen infection, or mechanical injury. There is no information on the ability of ROS to evoke systemic electrical or calcium signals in the model moss Physcomitrella and on the relationships between these responses. Here, we show that external application of hydrogen peroxide evokes electrical signals in the form of long-distance changes in the membrane potential, which transmit through the plant instantly after stimulation. The responses were calcium dependent, since their generation was inhibited by lanthanum, a calcium channel inhibitor (2 mM) or EDTA, a calcium chelator (0.5 mM). The electrical signals were partially dependent on glutamate receptor ion channels (GLR), since the knockout of GLR genes only slightly reduced the amplitude of the responses. The basal part of the gametophyte, which is rich in protonema cells, was the most sensitive to hydrogen peroxide. The measurements carried out on the protonema expressing fluorescent calcium biosensor GCaMP3 proved that. We also demonstrate upregulation of a stress-related gene which appears in a distant section of the moss 8 minutes after H_2_O_2_ treatment. The results help to understand the importance of both types of signals in the transmission of information about the appearance of ROS in the plant cell apoplast.

## Introduction

As an early terrestrial plant, the moss Physcomitrella (new botanical name *Physcomitrium patens*), from the beginning of the adaptation to land conditions, had to cope with exposure to UV radiation, ozone, wounding, and many other abiotic and biotic stressors that enhance the synthesis of reactive oxygen species (ROS) (Rensing et al., 2020, 2008; Tucker et al., 2005). Physcomitrella is an emerging model plant with a fully sequenced genome (Rensing et al., 2008; Lang et al., 2018). It is widely used to study defense mechanisms against environmental stress factors (Frank et al., 2005; Hoernstein et al., 2023; Koselski et al., 2019; Saidi et al., 2009). It was previously demonstrated in our lab that Physcomitrella is capable of generating action potentials (APs) after illumination with light of sufficient (over-threshold) intensity, cooling, glutamate (Glu) treatment, etc. (Koselski et al., 2020, 2019, 2008). Electrophysiological experiments with the application of ion channel inhibitors and manipulation of ion gradients across the plasma membrane indicated that Ca^2+^ fluxes from external and internal stores were involved in the generation of APs in Physcomitrella. They interacted with K^+^ and, to a lesser extent, with Cl^-^ fluxes. Our recent study on GCaMP mutants allowing monitoring of changes in the cytoplasmic Ca^2+^ concentration ([Ca^2+^]_cyt_) demonstrated that local Glu application caused an increase in [Ca^2+^]_cyt_ confined to the site of the Glu treatment, whereas AP was transmitted to distant cells (Koselski et al., 2020). This was rather an unexpected result, because in other plant species examined before, action potentials or variation potentials (VPs, known also as Slow Wave Potentials, SWPs) spread together with Ca^2+^ waves and seem mutually dependent (Choi et al., 2017, 2014; Gilroy et al., 2016). In the model vascular plant *Arabidopsis thaliana* two glutamate receptors *At*GLR3.3 and *At*GLR3.6 were identified as key players in wound-induced SWP generation and transmission (Farmer et al., 2020; Mousavi et al., 2013). Both genes are predominantly expressed in vascular tissues: phloem and xylem contact cells, respectively. Glutamate (and other amino acids) released from wounded tissues binds to the Ligand Binding Domain (LBD) subunit of the glutamate receptor-like channels facilitating opening of the channel pore and in consequence initiating a Ca^2+^ wave and other downstream responses, like jasmonate (JA) key genes and other stress-related gene expressions (Grenzi et al., 2022).

According to the recent findings, in vascular plants, in addition to the ion fluxes facilitating regenerative [Ca^2+^]_cyt_ wave formation and transmission, the ROS based component supplements the long-distance signaling system (Baxter et al., 2014; Gilroy et al., 2016; Kurusu et al., 2015; Marcec et al., 2019). NADPH oxidases (respiratory burst oxidase homologs, RBOHs) localized in the plasma membrane seem to be good candidates to fit this scheme. RBOHs cause reduction of oxygen to superoxide anion (O_2_^.-^), which is quickly converted to H_2_O_2_. They possess EF-hand motives which activate them upon binding of Ca^2+^ to enhance ROS production (Marcec et al., 2019; Wojtaszek, 1997). Before the machinery of ROS scavenging reduces the ROS level back to normal, its temporary excess affects many metabolic and signaling pathways and their components. H_2_O_2_ produced in the apoplast can be quickly transported to the cytosol via aquaporins (Tian et al., 2016). Recently, a specific apoplastic H_2_O_2_ receptor was discovered, Hydrogen-Peroxide-Induced Ca^2+^ Increases 1 (HPCA1) (Wu et al. 2020). It is a membrane-spanning protein composed of an extracellular H_2_O_2_ sensor and a cytoplasmic kinase component, which is postulated to activate Ca^2+^- permeable channels after H_2_O_2_ binding to the sensor (Foyer, 2020; Wu et al., 2020). Thus, ion channels can be affected from both sides of the plasma membrane. It was previously demonstrated that ROS can activate ion channels in different plant species classified in different branches of the phylogenetic tree of characean algae (Demidchik et al., 1997) to dicots and monocots (Wu et al., 2015).

In vascular plants, hydrogen peroxide and hydroxyl radicals affect Ca^2+^-permeable channels in the plasma membrane and in the internal store – the vacuole. In *Arabidopsis thaliana*, two classes of non-selective calcium-permeable channels are postulated to be involved: GLR channels activated by glutamate and cyclic nucleotide-gated channels CNGCs (Finka et al., 2012; Mousavi et al., 2013). Additionally, annexin1 has been reported to be involved in ROS-induced [Ca^2+^]_cyt_ elevation (Richards et al., 2014). It has been demonstrated that homologs of GLU and CNGC channels were involved in signaling processes in Physcomitrella (Finka and Goloubinoff, 2014; Koselski et al., 2020; Ortiz-Ramírez et al., 2017). In addition to calcium-permeable channels, K^+^ channels in the plasma membrane have been found to respond to an increase in the ROS level (Demidchik et al., 2010). Massive K^+^ efflux in ROS-treated plants was one of the first observations of plant response to this type of stress factors (Demidchik et al., 2010; Nassery, 1979). In Arabidopsis, GORK (Guard Cell Outward Rectifying K^+^) and SKOR (Stellar Potassium Outward Rectifier) channels have been identified to be responsible for that effect (Demidchik, 2018). These channels are also regarded as good candidates to pass an outward current during the AP repolarization phase in excitable plants (Cuin et al., 2018).

In the present study, we examined the effects of H_2_O_2_ treatment of intact Physcomitrella. Electrical potential changes and [Ca^2+^]_cyt_ transients were measured in wild type plants (WT) and *glr1^KO^* mutants treated with H_2_O_2_. We demonstrated that in the WT plants, H_2_O_2_ evoked APs recorded both in the gametophyte leaves and protonema. The susceptibility of the protonema to the treatment was much higher than in the leaf cells (0.5 mM versus 5 mM). These signals were blocked by La^3+^ - a calcium channel inhibitor and by EDTA – a Ca^2+^ chelator. In the *glr1^KO^*mutants, the electrical signals were reduced but not totally blocked, which reveals that GLR channels are not crucial in H_2_O_2_-induced membrane potential changes and Ca^2+^ fluxes in Physcomitrella. Involvement of other factors affecting [Ca^2+^]_cyt_ is discussed. Looking for downstream effects of the signals, we demonstrate an enhanced expression of stress-related genes.

## Results

### Electrical signals in leaf cells

The microelectrode measurements carried out on leaf (phylloid) cells from the Physcomitrella gametophyte proved that the application of hydrogen peroxide evoked systemic electrical signals in the form of membrane-potential changes transmitted along the plant. The long-distance responses transmitted from the basal part of the plant (from the rhizoid side) and recorded in apical leaf cells are presented in Fig. 1A. Experiments with different H_2_O_2_ concentrations indicate an influence of the concentration on amplitude and shape of membrane potential changes recorded during the response. Stimulation of the basal part of the plant with 0.05 mM and 0.1 mM H_2_O_2_ evoked irregular membrane potential changes with similar amplitude, duration, and rate of depolarization (Fig. 1A, Tab 1). After an increase of H_2_O_2_ concentration to 0.5 mM, significant changes in basic parameters describing the membrane potential changes were obtained (Fig. 1A, Tab 1). The cell membrane depolarized over twice as fast than in the lower concentrations (14.5±1.6 mV/s, n=18), and reached more positive values. The amplitude of the depolarization amounted to 68±3 mV (n=19) and was about 30 mV higher than after treatment with 0.05 and 0.1 mM, H_2_O_2_, respectively. Another difference occurred in the shape of the responses, where after the peak of depolarization evoked by 0.5 mM H_2_O_2_, a several-minute plateau of the membrane potential was recorded. Hydrogen peroxide in the concentration of 0.5 mM was used to compare transmission ability of electrical signals in the opposite direction - from the apical to the basal part of the plant (Fig. 1B). The electrical signals transmitted from the apical part of the plant characterized in a lower rate of depolarization (2.3±0.4 mV/s, n=8), amplitude (22±5 mV, n=9) and more negative values of maximum depolarization (-142±8 mV, n=9). The additional feature was an absence of a characteristic plateau of the membrane potential. The shape of the membrane potential changes resembled the responses obtained after direct stimulation of a cell in the apical part of the plant (Fig. 1C) or removing of the basal part (Fig. 1D), where a 10-fold increase in the H_2_O_2_ concentration (to 5 mM) evoked responses similar to these transmitted from the basal part after 0.5 mM H_2_O_2_ administration. These results indicate that the highest susceptibility to hydrogen peroxide occurs in the basal part of the plant having protonema cells and rhizoids, which act as an initial place for the generation of long-distance electrical signals propagated toward the plant apex. The results also showed that an increase of the H_2_O_2_ concentration facilitates generation of fully developed responses resembling action potentials (APs).

**Fig. 1.**
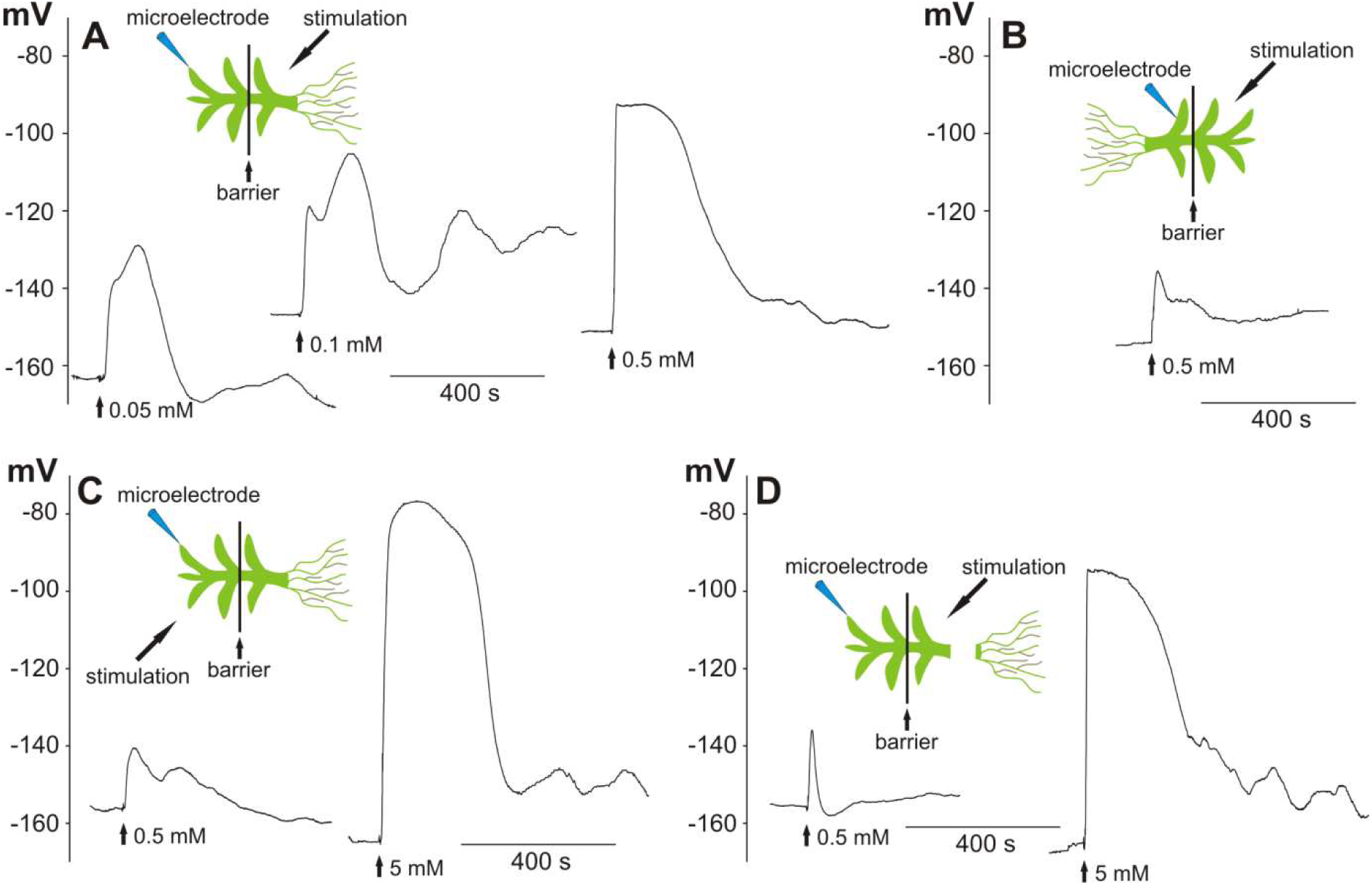
Hydrogen peroxide-evoked long-distance electrical signals recorded in a Physcomitrella leaf (phylloid) cell. The recordings were carried out in bipartite chambers with a barrier separating two parts of the gametophyte. Schemes presenting the site of stimulation and insertion of the microelectrode are in placed on the top of the recordings. A – long-distance electrical signals in the form of membrane potential changes recorded after the stimulation of the basal part of the gametophyte with different H_2_O_2_ concentrations. B - electrical signals recorded after stimulation of the apical part of the gametophyte. C and D - reduction of the sensitivity of the leaf cells to H_2_O_2_ recorded after direct stimulation of the tested cell located in the apical part of the gametophyte and in the gametophyte with a cut-off basal part, respectively. Vertical axes present the values of the membrane potential (in mV).

**Table 1.** Values of electrical signal parameters obtained in leaf cells after the application of hydrogen peroxide in different concentrations and different variants of experiments.

| | $V_0$ (mV) | $V_{\max}$ (mV) | A (mV) | $t_{1/2}$ (s) | $R_{\text{dep}}$ (mV/s) |
| --- | --- | --- | --- | --- | --- |
| **basal stimulation 0.5 mM** | -154±3<br>(n=19) | -86±3<sup>a, b, c, d, e</sup><br>(n=19) | 68±3<sup>a, b, c</sup><br>(n=19) | 236±44<sup>a</sup><br>(n=17) | 14.5±1.6<sup>a, b, c, d, e</sup><br>(n=18) |
| **basal stimulation 0.1 mM** | -161±4<br>(n=14) | -123±6<sup>d, i</sup><br>(n=14) | 38±6<sup>g</sup><br>(n=14) | 169±33<br>(n=13) | 6±1.4<sup>e</sup><br>(n=13) |
| **basal stimulation 0.05 mM** | -158±2<br>(n=12) | -121±7<sup>e, j</sup><br>(n=12) | 37±6<sup>h</sup><br>(n=12) | 140±15<br>(n=11) | 4.4±1<sup>d</sup><br>(n=11) |
| **apical stimulation 0.5 mM** | -165±5<br>(n=13) | -142±8<sup>a, f, k</sup><br>(n=9) | 22±5<sup>b, e, j</sup><br>(n=9) | 85±20<sup>c, f</sup><br>(n=8) | 2.3±0.4<sup>b</sup><br>(n=8) |
| **direct apical stimulation 0.5 mM** | -153±4<br>(n=9) | -136±7<sup>b, g, l</sup><br>(n=9) | 17±7<sup>a, d, i</sup><br>(n=9) | 89±28<sup>d, g</sup><br>(n=8) | 2.9±1.6<sup>a</sup><br>(n=6) |
| **direct apical stimulation 5 mM** | -153±5<br>(n=11) | -75±5<sup>f, g, h, i, j</sup><br>(n=11) | 78±4<sup>d, e, f, g, h</sup><br>(n=11) | 316±52<sup>e, f, g</sup><br>(n=11) | 11.6±3.5<br>(n=9) |
| **basal stimulation of plant without rhizoids 0.5 mM** | -156±3<br>(n=15) | -130±7<sup>c, h, m</sup><br>(n=12) | 24±5<sup>c, f, k</sup><br>(n=12) | 51±12<sup>a, b, e</sup><br>(n=12) | 3.3±0.8<sup>c</sup><br>(n=12) |
| **basal stimulation of plant without rhizoids 5 mM** | -153±3<br>(n=10) | -86±6<sup>k, l, m</sup><br>(n=10) | 67±6<sup>i, j, k</sup><br>(n=10) | 389±79<sup>b, c, d</sup><br>(n=9) | 10.5±2.1<br>(n=10) |

Action potentials belong to the basic long-distance electrical signals recorded in plant cells whose generation is dependent on an influx of calcium ions into the cytoplasm. We decided to study the dependence of the recorded responses on the presence of an inhibitor of calcium channels (2 mM lanthanum) or a calcium chelator (0.5 mM EDTA), respectively (Fig. 2 A, B). Apart from effects of lanthanum and EDTA, the possibility to reverse the evoked effects was also examined. Each plant was stimulated twice - after initial immersion for 3-4 hours in lanthanum or EDTA, and then after the exchange of the solution back to the standard solution. Immersion of the plants in lanthanum caused a total blockage of the response in 10 of the 19 tested cells. The responses in the other plants had a significantly reduced amplitude (to 9±2 mV, n=9). Lanthanum also shifted the resting potential to positive values (to -130±3 ± mV, n=19). The depolarization after the lanthanum application was slower (the rate amounted to 0.5±0.2 ±mV/s, n=8) and reached more negative values (-128±5, n=9). After a washout of lanthanum from the measuring chamber, the amplitude of the responses increased to 27±4 mV (n=4), which indicated that inhibition of the responses by lanthanum is partially reversible. EDTA totally blocked the responses in 9 of the 18 tested cells. Similar to lanthanum, EDTA evoked reduction of the amplitude (to 13±4 mV, n=9), shift of the resting potential to more positive values (to -122±5-± mV, n=18) and reduction of the rate of depolarization (to 1.4±0.7±mV/s, n=7). The shift of the resting potential after EDTA was partially reversible, since a washout resulted in a restoration of the resting potential close to the values measured in standard solution (-144±4 mV, n=18). The comparison of all these parameters of the membrane potential changes is compiled in Table 2.

**Fig. 2.**
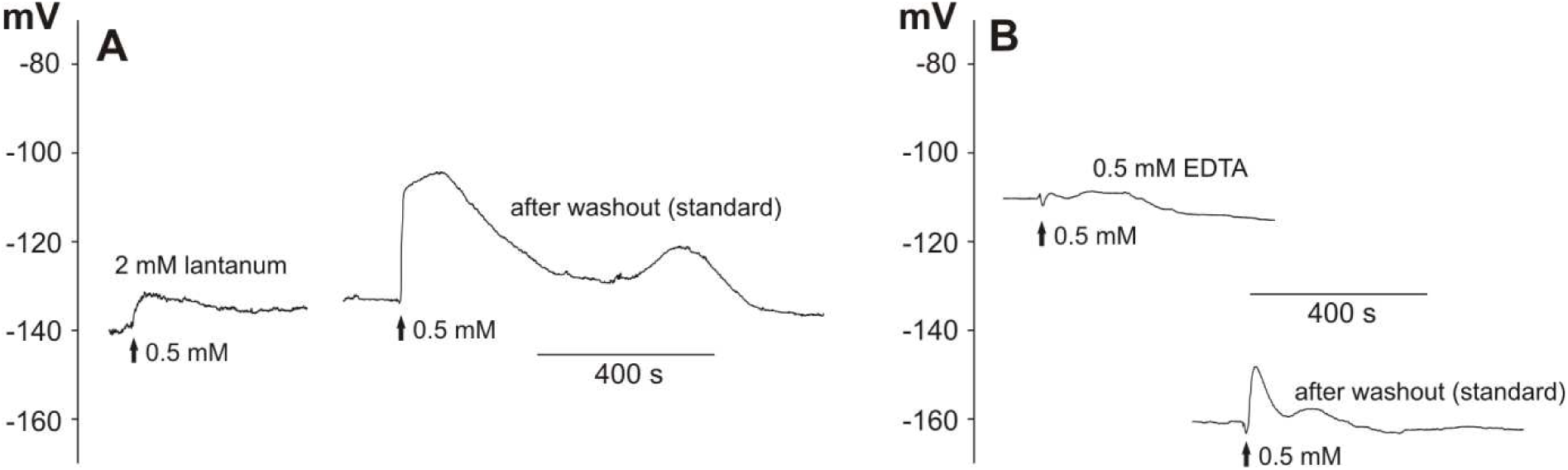
Blockage of H_2_O_2_-evoked long-distance electrical signals by 2 mM lanthanum (A) and 0.5 mM EDTA (B). The method of stimulation was the same as in Fig. 1A. The representative membrane potential changes show the effects of preincubation in lanthanum (A) or EDTA (B), respectively, and then washout of the drugs carried out on the same plant.

**Table 2.** Values of electrical signal parameters obtained in leaf cells after the application of 0.5 mM hydrogen peroxide in the basal part of the gametophyte.

|  | **V<sub>0</sub> (mV)** | **V<sub>max</sub> (mV)** | **A (mV)** | **t<sub>1/2</sub> (s)** | **R<sub>dep</sub> (mV/s)** |
| --- | --- | --- | --- | --- | --- |
| **standard (WT)** | -154±3<sup>a, b, c</sup><br>(n=19) | -86±3<sup>a, b, c</sup><br>(n=19) | 68±3<sup>a, b, c, d</sup><br>(n=19) | 236±44<br>(n=17) | 14.5±1.6<sup>a, b, c, d</sup><br>(n=18) |
| **2 mM La<sup>3+</sup>** | -130±3<sup>b</sup><br>(n=19) | -128±5<sup>a, d</sup><br>(n=9) | 9±2<sup>a, e</sup><br>(n=9) | 205±25<br>(n=3) | 0.5±0.2<sup>a, e</sup><br>(n=8) |
| **after washout of 2 mM La<sup>3+</sup>** | -130±4<sup>c</sup><br>(n=19) | -104±5<br>(n=5) | 27±4<sup>d</sup><br>(n=4) | 224±34<br>(n=15) | 3.6±0.8<sup>c</sup><br>(n=14) |
| **0.5 mM EDTA** | -122±5<sup>a, d, e</sup><br>(n=18) | -115±7<sup>c</sup><br>(n=9) | 13±4<sup>b</sup><br>(n=9) | 126±32<br>(n=6) | 1.4±0.7<sup>b, f</sup><br>(n=7) |
| **after washout of 0.5 mM** | -144±4<sup>e</sup><br>(n=18) | -121±7<sup>b</sup><br>(n=8) | 24±6<sup>c</sup><br>(n=8) | 63±14<br>(n=7) | 2.5±0.7<sup>d</sup><br>(n=7) |
| **standard (Pp*glr1*<sup>KO</sup>)** | -146±3<sup>d</sup><br>(n=26) | -100±4<sup>d</sup><br>(n=26) | 46±3<sup>e</sup><br>(n=26) | 194±21<br>(n=25) | 9.6±1.2<sup>e, f</sup><br>(n=24) |

### Electrical signals in protonema cells

The results of the measurements carried out on the leaf cells indicated that it is hard to evoke electrical signals in such cells, and the responses can start mainly from the protonema cells and/or rhizoids - probably the target for the action of hydrogen peroxide. In order to study this hypothesis, we decided to examine the effect of H_2_O_2_ on electrical membrane potential changes in protonema cells. In these measurements, stimulation was carried out by microinjection of H_2_O_2_ in three regions: initially into a chain of protonema cells adjacent to the tested cell with the inserted microelectrode, then directly into the tested cell, and at the end into the gametophyte base.

The results confirmed that, although electrical signals in the chain of protonema cells are transmitted from cell to cell, the responses recorded in the same cell differed depending on the region of stimulation (Fig. 4, Tab. 3, Supplementary Video S1). The stimulation of the gametophyte base was the most effective, as it evoked electrical signals in each tested cell immediately upon stimulation. In comparison to the electrical signals recorded in the leaf cells, the signals from the protonema reached a higher amplitude (81±4 mV, n=8) and duration (486±100 s, n=8). Surprisingly, weaker effects were achieved by the direct stimulation of the tested cell or the stimulation of adjacent cells located close to the tested cell. This method of stimulation evoked electrical signals with a smaller amplitude (66±6 mV, n=8) than when H_2_O_2_ was applied in the gametophyte base. The responses recorded in the protonema after the direct or indirect stimulation also exhibited a low depolarization rate (1.6±0.6 mV/s, n=8), which was lower than the responses to the stimulation of the basal part of the gametophyte (9.3±2.5 mV/s, n=8) or than those recorded in the leaf cells. Cell-to-cell transmission of the electrical signal evoked by stimulation of the single cells from the chain of protonema cells was rarely recorded and occurred only in 2 out of 18 tested plants (Supplementary Fig. 1 and Video 2).

**Table 3.**
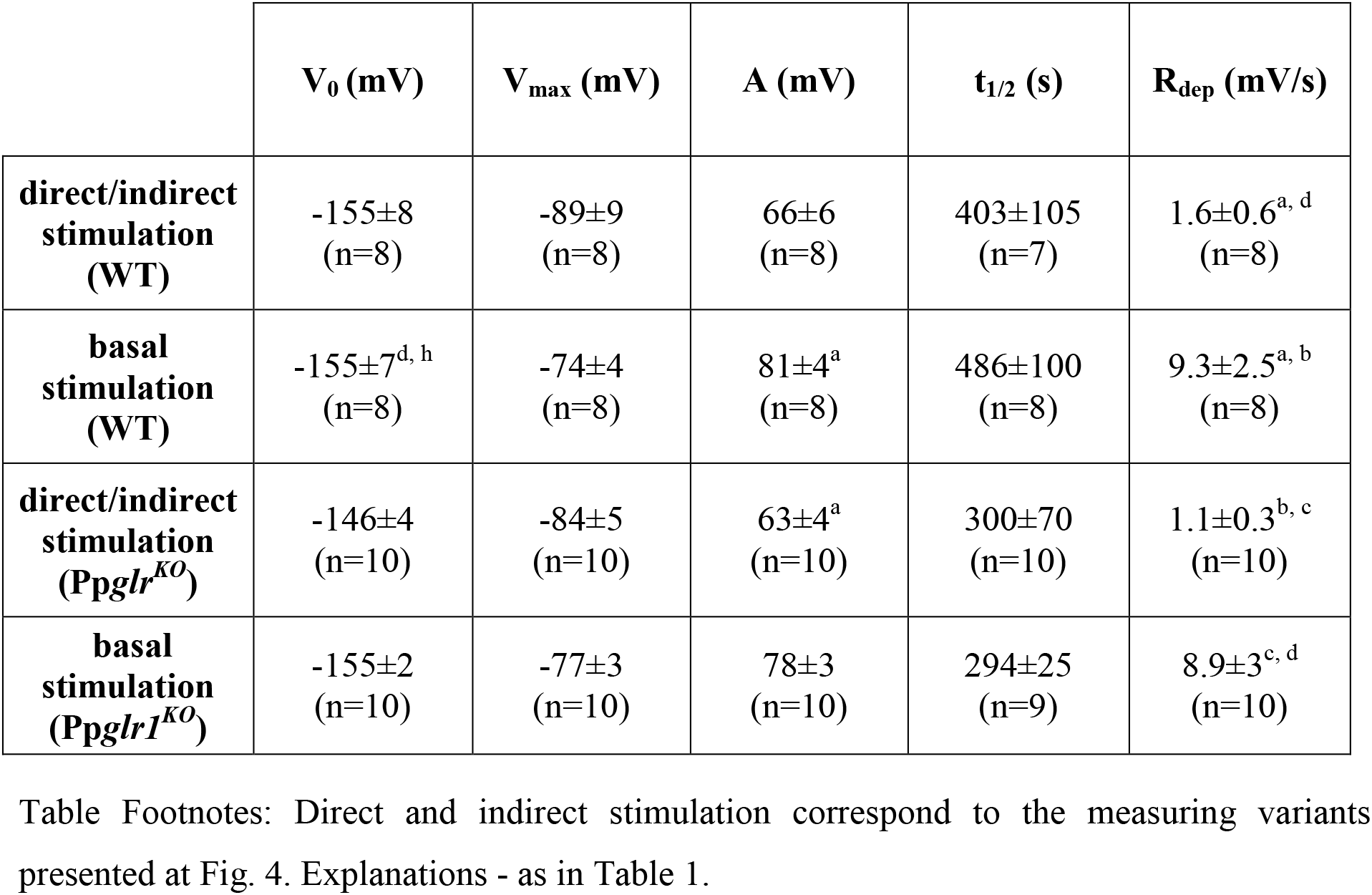
Values of parameters of electrical signals in protonema cells after the application of 0.5 mM hydrogen peroxide directly/indirectly to the cell or in the basal part of the gametophyte.

One of the candidates responsible for cell-to-cell communication in plants is the GLR receptor, which can act as a non-selective calcium-permeable channel. The participation of the GLR receptor in the transmission of electrical signals recorded in our experiments was tested with the use of a *glr1* mutant of Physcomitrella (Pp*glr^KO^*). In the Pp*glr1^KO^*mutants, as in the wild type (WT), electrical signals propagated from the base of the gametophyte to the leaf and protonema cells (Fig. 3, 4, Supplementary Video S3). In comparison to WT, electrical signals recorded in the leaf cells of Pp*glr1^KO^* had a reduced amplitude (to 46±3 mV, n=26) and a lower depolarization rate (9.6±1.2 mV/s, n=24). The amplitude and rate of depolarization of responses in the protonema cells from Pp*glr1^KO^*recorded after the stimulation of the basal gametophyte part reached similar values to those recorded in the WT protonema cells (78±3 mV, n=10 and 8.9±3 mV/s, n=10, respectively). As in the WT protonema cells, the direct or indirect stimulation of the protonema cells from Pp*glr1^KO^* evoked a smaller amplitude (63±4 mV, n=10) and a lower depolarization rate (1.1±0.3 mV/s, n=10) than after the stimulation of the basal part of the Pp*glr1^KO^*gametophyte. All these changes in the parameters of changes in membrane potential in the protonema cells are presented in Table 3.

**Fig. 3.**
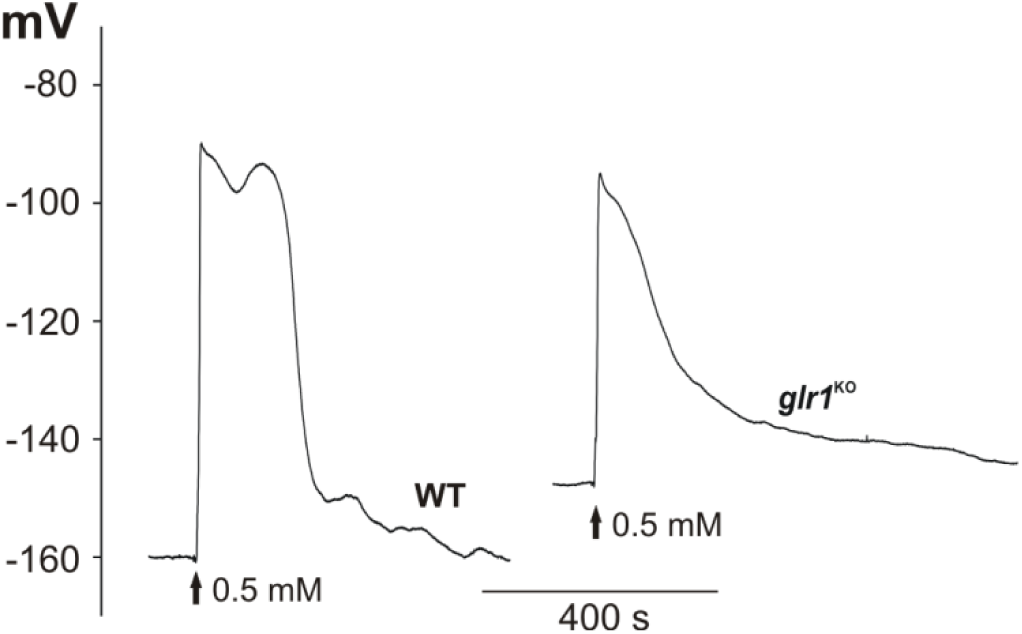
Comparison of H_2_O_2_-evoked long-distance electrical signals recorded in wild type and Pp*glr1^KO^* mutant. The method of stimulation was the same as in Fig. 1A.

**Fig. 4.**
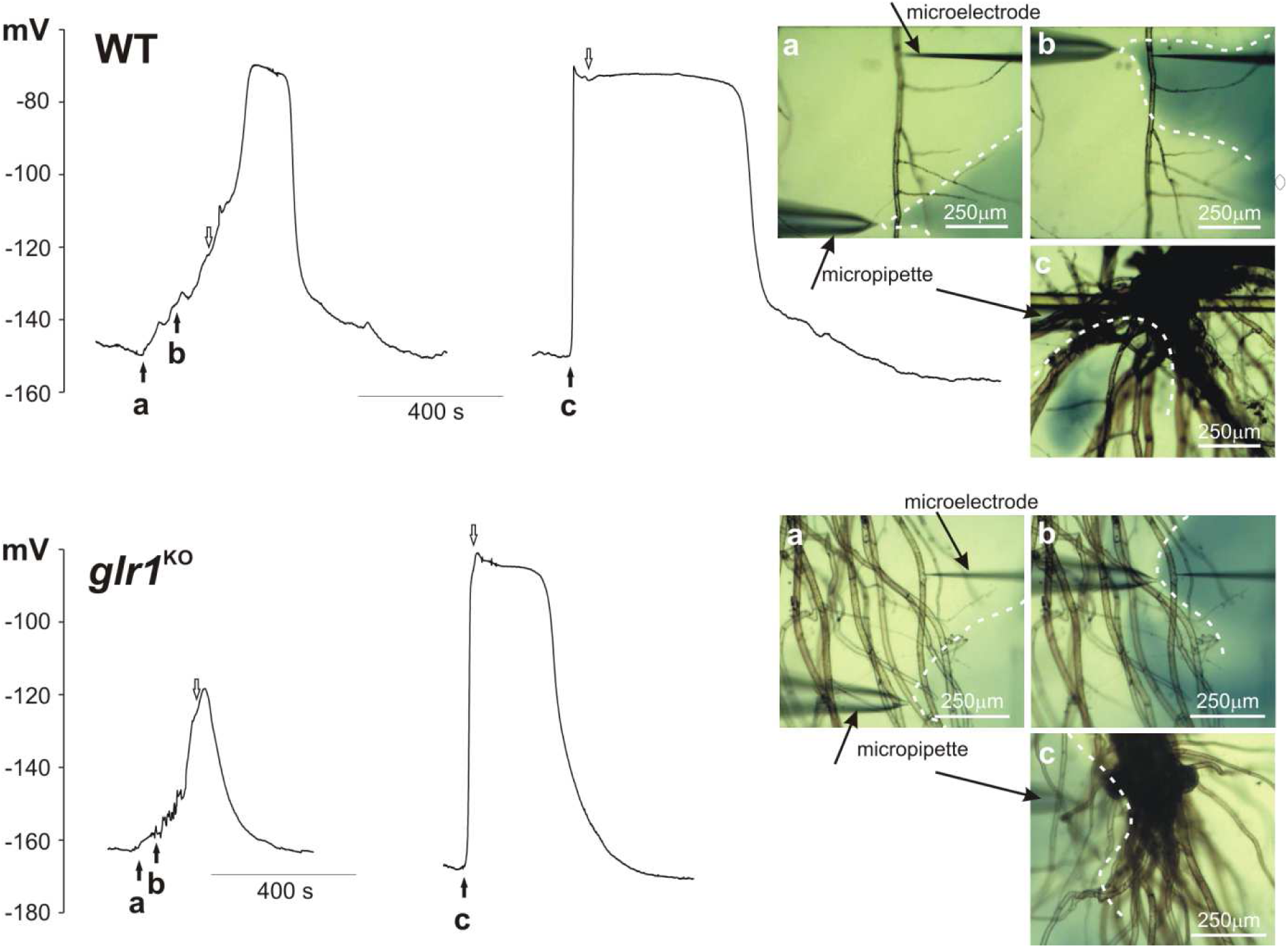
Hydrogen peroxide-evoked membrane potential changes recorded in protonema cells from wild type and Pp*glr1*^KO^ mutant. Hydrogen peroxide (0.5 mM) was injected into the selected site by the micropipette. In the pictures placed on the right side of presented traces, three different measurement variants are presented: a - stimulation of the cells adjacent to the tested cell (site of the microelectrode insertion), b - direct stimulation of the tested cell, and c - stimulation of the basal part of the gametophyte. The pictures show dispersion of H_2_O_2_ stained with 1 mM aniline blue (marked by dashed white lines) recorded 5 seconds after time points a, b, and c marked on the trace by black arrows. The end of stimulation (removal of the micropipette from the measuring chamber) was marked by white arrows.

### Calcium signals

Calcium signals were recorded in plants expressing GCaMP3, i.e. a fluorescent calcium indicator. Fluorescence measurements of calcium signals recorded after the stimulation of the basal part of the plant indicated that the application of 0.5 mM H_2_O_2_ evoked calcium waves which propagated from cell to cell in thread-like protonema cells. In contrast to electrical signals, the calcium signals were slower and appeared a few minutes after the application of the stimulus (Fig. 5, Supplementary Video S4). In some plants, the calcium signals did not start at the stimulation site but appeared at some distance from the stimulation, acting as a new source for calcium waves. In such a situation, calcium wave propagation was observed in both directions - into and from the stimulation site. The rate of calcium signals propagation along the protonema towards the site of the stimulation was similar to that in the opposite direction, reaching 5.5±0.7 µm/s (n=4) and 5.2±0.3 µm/s (n=7), respectively. A characteristic trait of some recorded calcium signals was its decrement with distance (Supplementary Fig. 2, Supplementary Video S5). As in the case of electrical signals, the observation of calcium signals in the stimulated leaves was possible after the increase of the hydrogen peroxide concentration to 5 mM (Supplementary Video S6). In turn, in contrast to electrical signals, calcium signals in leaf cells after application of 0.5 mM H_2_O_2_ to the basal part of the plant were observed occasionally only in several cells from single leaves, but never (in none of 30 tested plants) in all leaves (Supplementary Video S7).

**Fig. 5.**
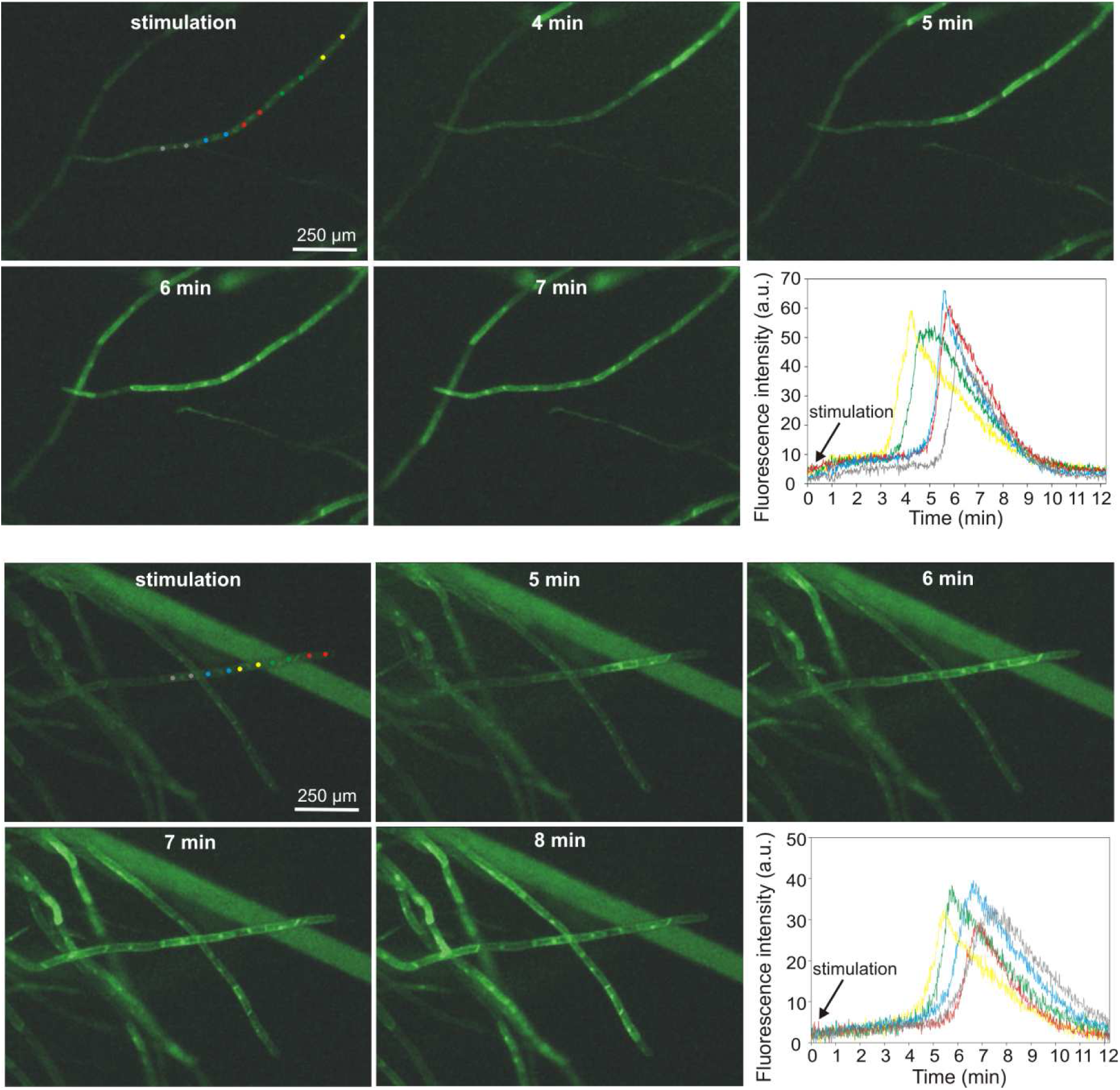
Hydrogen peroxide-evoked calcium signals recorded in protonema cells expressing fluorescent calcium biosensor GCaMP3. For each cell two circular ROIs (Region of Interest) located in cytoplasmic region from both sides of the nucleus were analyzed. The two panels of pictures show two different types of calcium signal propagation - at the upper the signal propagate in one direction starting from the site of stimulation (basal part of the gametophyte) and at lower panel, the signal propagation starts from one cell (marked by yellow ROIs) and propagate in two opposite directions. Recorded during the experiment changes in fluorescence intensity in different ROIs are presented in the bottom right corner of each picture panels.

### Expression of genes in distant regions of the moss

To analyze whether the electrical signal which was triggered upon local treatment would be accompanied by changes in gene expressions in distant tissues, we selected candidate genes and performed quantitative real-time PCR (qPCR). The candidates were selected based on homology to Arabidopsis genes differentially expressed after local stress stimulus (Zandalinas et al., 2019). In that study, a high light stimulus was locally imposed on selected Arabidopsis rosette leaves and changes in gene expression in the locally treated and the distant, non-treated leaves (local and systemic response, respectively) were investigated by RNAseq analysis. From this published list, we selected candidate genes which were upregulated in the systemic response if their expression was also inducible via H_2_O_2_ treatment and depended on the function of the respiratory burst oxidase homolog D (RBOHD). Here, the gene encoding a galacturonosyltransferase-like 10 protein (AT3G28340) was 5-fold upregulated 5 minutes after the light stress treatment (Zandalinas et al., 2019). In Physcomitrella, we identified four homologous proteins (Pp3c5_28420V3.1, Pp3c25_14930V3.1, Pp3c16_25090V3.1, Pp3c2_18670V3.1) via *BlastP* search (Altschul et al., 1997) against the Physcomitrella proteome (Lang et al., 2018). In analogy to Zandalinas et al. (2019) we tested their responsiveness to H_2_O_2_ treatment and submerged entire gametophores in 0.5 mM H_2_O_2_ for 8 minutes. Here, we employed hydroponic gametophore cultures and analyzed the expression of the candidate genes with qPCR. Of the four Physcomitrella homologues, only one gene (Supplementary Fig S3, Pp3c16_25090V3.1) exhibited a significant increase in gene expression (p = 0.0305) and hence, was selected for further analysis. For Pp3c5_28420V3.1. an increase in gene expression was detectable (Supplementary Fig S3) but the difference was not significant. Nevertheless, this gene was included for further experiments.

For these two selected candidates we tested whether both genes are regulated in gametophore sections distant from a local treatment with H_2_O_2_. Apices of gametophores were harvested after treatment with 0.5 mM H_2_O_2_ at the base for 8 min (Supplementary Fig S4) as well as corresponding untreated apices. A significant upregulation of Pp3c16_25090V3.1 (p = 0.00184) was observed (Fig. 6) whereas an upregulation of Pp3c5_28420V3.1 was indicated, but was not statistically significant (p = 0.06075). Thus, a local stress stimulus (here: H_2_O_2_ at the gametophore base) triggers alteration of gene expression within 8 minutes in distant sections (here: apex) in Physcomitrella.

**Fig. 6.**
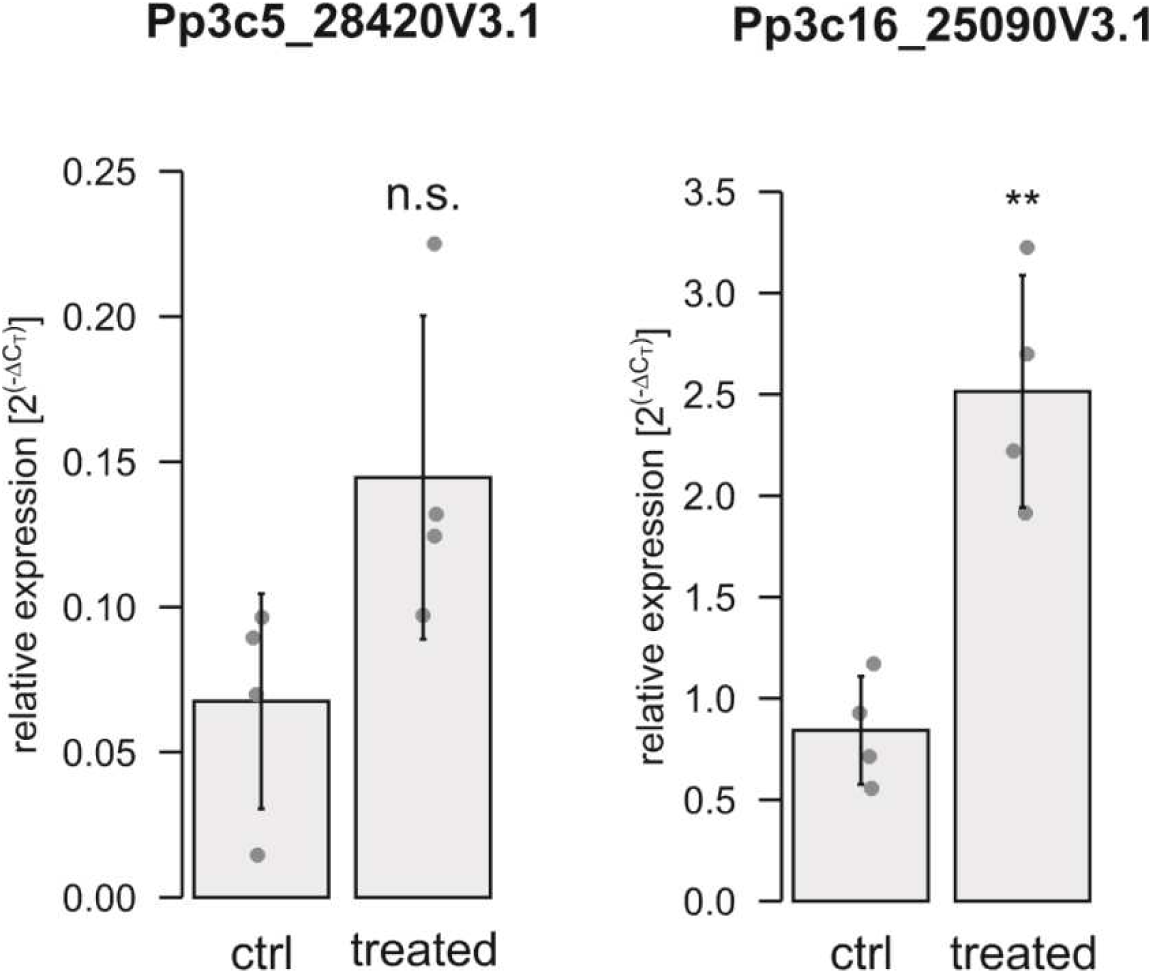
Gene expression analysis by qPCR of selected gene candidates. Apices were used from untreated gametophores and gametophores treated at the base with 0.5 mM H_2_O_2_ for 8 min. Dots represent biological replicates (mean values from three technical replicates). Bars are mean values from the four biological replicates with standard deviation. Relative expression (2^(-ΔCT)^) of Pp3c5_28420V3.1 and Pp3c16_25090V3.1 is calculated against the reference genes L21 (Pp3c13_2360V3.1) and LWD (Pp3c22_18860V3.1) according to Livak & Schmittgen (2001). Significance levels are based on a one-way Anova with subsequent post-hoc test (**p < 0.01); n.s. = not significant.

## Discussion

Plant cell signaling is based mainly on generation and transmission of different types of signals including electrical, calcium, and reactive oxygen species (ROS). Investigations of the relationship between these signals allow unraveling signaling pathways triggered in response to biotic and abiotic stresses. In this study, we tried to answer the question whether external application of ROS (hydrogen peroxide) evokes systemic response in the form of electrical and calcium signals, to analyze the interdependence of these signals and their impact on gene expression in distant regions.

The importance of ROS in the plant defense system has been evidenced by many experiments (Huang et al., 2019). One of the first studies focused on the involvement of ROS in cell-to-cell communication in *Arabidopsis* (Miller et al., 2009) proved that local stimuli produced ROS waves propagating through the plant at a rate of 8.4 cm/min and the response was dependent on a gene RBOHD (Respiratory Burst Oxidase Homolog D) encoding plant NADPH oxidase involved in the production of ROS. The rate of propagation of such RBOHD-related ROS signals recorded in different tissues ranged from ∼400 to 1400 µm/sec and was dependent on the type of stress (Choi et al., 2017). Together with propagation of the signal, the accumulation of extracellular ROS was observed along the path of the signal (Miller et al., 2009), indicating that each cell along the path is able to activate RBOHD and release ROS, which in turn trigger adjacent cells to carry out the same process. In such an autopropagation process, named ROS-induced ROS-release (RIRR), hydrogen peroxide generated by RBOHD can be regarded as a long-distance signal. However, in the study by Miller et al. (2009), external application of H_2_O_2_ did not evoke a ROS wave indicating involvement of some other signal molecules in the propagation of the ROS wave.

Calcium is a candidate for such a signal molecule and together with ROS can cooperate in long-distance transmission of information about stimuli. For example, calcium-dependent protein kinase CPK5 playing a role in plant immunity is important for cell-to-cell communication based on ROS waves (Dubiella et al., 2013). There are also other ways of activation of RBOH proteins by calcium, including direct binding of this ion to the EF-hand motif on the RBOH protein (Kimura et al., 2012) or binding of phosphatidic acid to the same protein whose accumulation in the cell can be induced by calcium (Zhang et al., 2009). In addition to the calcium-induced ROS release process, ROS-induced calcium release is possible. For example, it has been shown that ROS can activate different calcium permeable channels, e.g. hyperpolarization-activated Ca^2+^ channels in root cells (Demidchik et al., 2007), Ca^2+^ influx channels in guard cells (Pei et al., 2000), or Ca^2+^ permeable channels regulated by annexin1 (Richards et al., 2014). Recent studies carried out on Arabidopsis allowed the discovery of a plasma-membrane leucine-rich-repeat receptor kinase, i.e. hydrogen-peroxide induced Ca^2+^ increase 1 (HPCA1), which links the perception of apoplastic H_2_O_2_ with Ca^2+^ signaling (Wu et al., 2020). One of the most probable mechanisms of the cooperation between calcium and ROS in long-distance transmission of signals is that an increase in cytoplasmic calcium evoked by a local stimulus can induce production of ROS via Ca^2+^-RBOH interactions and accumulation of ROS in the apoplast. The transport of ROS from the apoplast to the cell, probably carried out by aquaporins or other channels, would evoke ROS-induced calcium release, which in turn could induce ROS production (Gilroy et al., 2014).

As shown by the results of our study, H_2_O_2_-evoked calcium signals are not systemic responses, such as those observed earlier in *P. patens* under osmotic stress and salt stimulation (Storti et al., 2018). Calcium waves with a velocity of about 400 µm/s in response to local salt stress were also measured in Arabidopsis (Choi et al., 2014), where blockage of calcium channels by lanthanum inhibited not only the calcium waves but also the ROS-regulated transcriptional marker (ZAT12), indicating that calcium and ROS waves are closely linked. Assuming the interconnection of calcium signals and ROS waves, it is unlikely that the extracellular application of H_2_O_2_ in our experiments evoked the ROS wave, since the H_2_O_2_-evoked calcium signals admittedly appeared at some distance from the stimulation site but propagated with a decrement (Fig. 3). The other feature of the H_2_O_2_-evoked calcium signals in Physcomitrella, i.e. the slower rate of propagation than in the case of the ROS wave (above 5 µm/s), also indicates a low probability of appearance of a calcium-associated ROS wave. The ability to evoke local calcium signals but not self-propagating calcium waves by the extracellular H_2_O_2_ application implies engagement of some other factors taking part in the transmission of information about such stimuli. It seems that, in *P. patens*, electrical signals, which appear at the moment of stimulation and propagate along the whole plant, are a proper carrier of the information about H_2_O_2_ enhancement. There still remains the question of the relationships between electrical, calcium, and ROS signals.

The relationship between ROS and electrical signals was previously confirmed in Arabidopsis mutants lacking RBOHD, where propagating electrical signals after heat stress or high light had a significantly reduced amplitude or were totally blocked in comparison to the wild type (Suzuki et al., 2013). Taking into account the ability of extracellular ROS (mainly hydrogen peroxide) to activate different ion channels (Demidchik, 2018), including calcium-permeable channels which can participate in long-distance electrical signals, it seems that ROS can promote electrical signals along the ROS wave path. This hypothesis can be also supported by the similar velocity of different types of plant self-propagating electrical signals to the velocity of the ROS wave [like action potentials (20-400 cm/min) and system potentials (5-10 cm/min); (Zimmermann et al., 2009)]. In Physcomitrella, the H_2_O_2_-evoked electrical signals propagated immediately from the basal part along the whole plant (Fig, 1 and 2), indicating that the signals appeared before the slowly propagating calcium waves. In cooperation with ROS and calcium waves, electrical signals, which are probably the fastest carriers of information, can initiate the whole array of plant defense responses. It is also probable that, by reaching cells distant from the stimulation site, electrical signals initiate a calcium-wave response only in some cells, in which the threshold of calcium signal generation is lower than in other cells. The relevance of this assumption is proved by the features of some calcium waves observed in the protonema cells, where at a distance from the stimulation site in the basal part of the plant, only a few protonema cells in the thread-like chain generated calcium signals propagating in two directions - into and out of the site of stimulation.

One of the candidates of ion channels participating in long-distance electrical signals in plants are glutamate receptor-like channels (GLR), which was confirmed in experiments carried out on Arabidopsis (Mousavi et al., 2013; Salvador-Recatalà, 2016; Salvador-Recatalà et al., 2014). Wound-induced electrical signals recorded in this species were dependent on clade III GLRs (GLR 3.3 and GLR 3.6) and played a crucial role in the distal production of jasmonates taking part in plant defense responses (Mousavi et al., 2013). It is a matter of discussion if those channels are directly responsible for Ca^2+^ fluxes since they are predominantly expressed in endomembranes (Farmer et al., 2020; Nguyen et al., 2018).

In Physcomitrella, only two *GLR* genes have been identified (Verret et al., 2010). Both genes (*PpGLR1* and *PpGLR2*) are paralogs to *GLR* clade III from Arabidopsis (De Bortoli et al., 2016) and encode channels participating in chemotaxis and reproduction of Physcomitrella (Ortiz-Ramírez et al., 2017). A patch-clamp study indicated that GLR1 is a calcium-permeable ion channel localized in the cell membrane and partially inhibited by glutamate receptor antagonists (Ortiz-Ramírez et al., 2017). These data indicate that GLR1 in Physcomitrella, similar to GLR 3.3 and GLR 3.6 from Arabidopsis, can be important in the transmission of long-distance electrical signals. This hypothesis was not fully confirmed in our study, since H_2_O_2_ evoked long-distance electrical signals in the Physcomitrella *glr1^KO^* mutants; however, the response differed in the amplitude (Tab. 1). Such a "weak" effect of the GLR1 receptor knockout suggests that, in addition to GLR channels, other channels must be engaged in the long-distance propagation of ROS-induced electrical signals. The most important channels taking part in the generation of electrical signals in Physcomitrella are probably calcium-permeable, given the blockage of H_2_O_2_-evoked responses by the calcium channel inhibitor (lanthanum) or the calcium chelator (EDTA) (Fig. 1D). It is also probable that, in addition to ion channels in the plasma membrane, a significant role in the transmission of electrical signals is assigned to intracellular calcium channels. One of the best-known intracellular channels permeable e.g. to calcium is the two-pore channel 1 (TPC1) located in the vacuolar membrane, the tonoplast (Dadacz-Narloch et al., 2011). A patch-clamp study carried out in our laboratory demonstrated that TPC channels in Physcomitrella vacuoles conduct Ca^2+^ currents (Koselski et al., 2013). Involvement of the channel in the transmission of long-distance calcium waves has been demonstrated (Choi et al., 2014), but there is no information about the role of TPC1 in the transmission of electrical signals. As calcium-permeable channels, TPC1 may be involved in the release of calcium from intracellular compartments (Qudeimat et al., 2008) leading to calcium-induced calcium release (CICR), a desired phenomenon for long-distance signal transmission, but this assumption arouses controversy (Pottosin et al., 1997; Ward et al., 1994). The role of H_2_O_2_ in CICR is questionable since MIFE (Microelectrode Ion Flux Estimation) and patch-clamp studies carried out on vacuoles from *Beta vulgaris* demonstrated that H_2_O_2_ suppressed the Ca^2+^ efflux from the vacuole and slow vacuolar (SV) currents carried by TPC1 (Pottosin et al., 2009).

Effectively, the treatment with 0.5 mM H_2_O_2_ at the base of gametophores was sufficient to trigger an increase in gene expression in the apex of a component of the homogalacturonan biosynthesis (Fig. 6, Pp3c16_25090V3.1). Here, the homologous candidate Pp3c5_28420V3.1 was not significantly upregulated although an increasing trend was detectable. This agrees with publicly available data indicating that both genes are not co-expressed. Galacturonosyltransferase-like proteins such as Pp3c16_25090V3.1 act in the pectin assembly (homogalacturonan biosynthesis) of the primary cell wall (reviewed in Loix et al., 2017). Under oxidative stress, cell-wall pectins also represent a source for the biosynthesis of ascorbic acid (García-Caparrós et al., 2021; Valpuesta and Botella, 2004), which in turn is used to detoxify ROS such as H_2_O_2_. Both selected candidates are homologues of an Arabidopsis isoform (AT3G28340) whose gene expression is regulated via ROS waves (Zandalinas et al., 2019). However, it should be noted that Galacturonosyltransferase-like proteins comprise a large gene family with at least 21 members in Arabidopsis and 17 genes in Physcomitrella (Van Bel et al., 2022) and clear ortholog relations are not yet resolved. The expression of the Arabidopsis homolog (AT3G28340) was 5-fold increased only in systemic leaves distant from a local high light stress impulse (Zandalinas et al., 2019) and those data further indicate that the increase of expression was a response to a ROS wave. In contrast, the two selected homologues in Physcomitrella were not regulated by high light stress (Supplementary Fig S5) but their expression increased after heat stress. Heat stress in plants is accompanied by the elevated production of reactive oxygen species (ROS) such as H_2_O_2_ (reviewed in Mittler et al., 2022). In summary, these data show that the gene expression of at least one of the two Physcomitrella candidates (Pp3c16_25090V3.1) is responsive to ROS. Consequently, the increase of expression in the untreated apex (Fig. 6) was likely based on a propagating ROS-wave.

Taken together, our study demonstrates differences in the generation and propagation of H_2_O_2_-evoked electrical and calcium signals in the model moss Physcomitrella. Many of the applied variants of measurements indicated that the basal part of the gametophyte is the most excitable region, probably because a large number of protonema cells are juvenile cells from an early stage of new gametophyte development. In comparison to the leaf cells, the responses in the protonema exhibited higher amplitudes and lasted longer, which may indicate higher susceptibility of such cells to ROS (Table 2 and 3, Fig. 1 and 4). The main difference between the electrical and calcium signals was the velocity of propagation, which was higher in the electrical signals. The other difference was the ability to propagate without a decrement; in contrast to the electrical signals, calcium signals diminished with distance, even if they appeared in some distance from stimuli (Fig. 5, Supplementary Video S4 and S5). Given these differences, we propose that H_2_O_2_-evoked long-distance electrical signals are the first to reach distant regions of the plant and activate calcium signals, but not in every cell. The protonema cells were the most susceptible to calcium signals, indicating that the signals play a key role in young and developing cells. A similar observation was reported in our previous work focused on glutamate-evoked calcium signals (Koselski et al., 2020). Electrical and calcium signals were not the only effect of H_2_O_2_ application. Gene expression analysis proved that apart from the signals, an increase of stress-related genes expression is observed in not stimulated distant regions of plants (Fig. 6). The increase in the gene expression appeared after 8 minutes - the time close to duration of electrical signals. Similar time scale of electrical signals and genes expression, raises the question about interdependence of the both phenomena.

## Materials and methods

### Cultivation of plant material used for electrophysiological and calcium imaging analysis

Physcomitrella gametophytes were grown on KNOP solid agar medium (Reski and Abel, 1985) in 160 mm diameter Petri dishes. The wild-type plants (WT) and knockout *glr1* mutants (Pp*glr1*^KO^) were used for the analysis of membrane potential changes in leaf and protonema cells with use of the microelectrode technique. A Physcomitrella mutant expressing GCaMP3 was used for the fluorescence imaging of changes in the calcium concentration. The plants were grown in a growing chamber (Conviron Adaptis A1000, Conviron, Winnipeg, Canada) at a photoperiod of 16/8 light/dark, with light intensity 50 µmol/m^2^ s, and at temperature set to 23°C.

### Cultivation of plant material used for gene expression analysis

Physcomitrella WT protonema (new species name: *Physcomitrium patens* (Hedw.) Mitt. (Medina et al., 2019); ecotype “Gransden 2004” was cultivated in Knop medium with microelements. Knop medium (pH 5.8) containing 250 mg/L KH_2_PO_4_, 250 mg/L KCl, 250 mg/L MgSO_4_ x 7 H_2_O, 1,000 mg/L Ca(NO_3_)_2_ x 4 H_2_O and 12.5 mg/L FeSO_4_ x 7 H_2_O was prepared as described (Reski and Abel, 1985) and 10 mL per litre of a microelement (ME) stock solution (309 mg/L H_3_BO_3_, 845 mg/L MnSO_4_ x 1 H_2_O, 431 mg/L ZnSO_4_ x 7 H_2_O, 41.5 mg/L KI, 12.1 mg/L Na_2_MoO_4_ x 2 H_2_O, 1.25 mg/L CoSO_4_ x 5 H_2_O, 1.46 mg/L Co(NO_3_)_2_ x 6 H_2_O) as described (Egener et al., 2002; Schween et al., 2003). The suspension culture was dispersed weekly with an ULTRA-TURRAX (IKA) at 18,000 rpm for 90 s.

Hydroponic cultures of Physcomitrella gametophores were cultivated as described in Hoernstein et al. (2023). Glass rings covered with mesh (PP, 250 m mesh, 215 m thread, Zitt Thoma GmbH, Freiburg, Germany) were prepared as described in Erxleben et al. (2012). Protonema suspension was adjusted to a final density of 440 mg/L (dry weight per volume) as described (Decker et al., 2017) and evenly distributed in equal volumes on the mesh surface. Glass rings covered with protonema were placed in Magenta^®^Vessels (Sigma-Aldrich, St. Louis, USA). KnopME medium was supplemented until it touched the bottom of the mesh. The medium was changed every 4 weeks.

Suspension cultures and hydroponic cultures were cultivated under standard light conditions (55 µmol photons/m^2^s) at 22°C in a 16h/8h light/dark cycle.

### Measurements of membrane potential in leaf cells

The method of membrane-potential measurements with microelectrodes was similar to that employed in Koselski et al. (2020). Plastic Petri dishes used in the microelectrode measurements were divided into two chambers with a barrier. The barrier had a small (1 mm width) gap sealed with Vaseline. Before the experiments the plants were incubated for 3-6 hours at light intensity of 50 µmol/m^2^ in a bath solution containing (in mM) 1 KCl, 1 CaCl_2_, 50 sorbitol, and 2 HEPES, pH 7.5 (buffered by Tris). The stimuli were introduced by application of 500 µL of a bath solution supplemented with 0.5 mM hydrogen peroxide. The lanthanum chloride or EDTA influence on the H_2_O_2_-evoked responses described in this study was assessed by application of one of these substances into one of the Petri dish compartments containing the basal part of the plant. Borosilicate glass capillaries (1B150F-6, World Precision Instruments, Sarasota, USA) were used to make micropipettes with the use of a P-30 micropipette puller (Shutter Instrument Co., Novato, USA), filled with 100 mM KCl, and connected to the FD223 electrometer (World Precision Instruments, Sarasota, USA). A Sensapex SMX (Sensapex, Oulu, Finland) electronic micromanipulator was used for positioning and insertion of the microelectrode. The reference electrode was composed of an Ag/AgCl_2_ wire inside a plastic tube filled with 100 mM KCl and ending with a porous tip. The measured data were acquired by a Lab-Trax-4 device (World Precision Instruments, Sarasota, USA) working with LabScribe2 software, which also allowed analyses of the data. The analysis of statistical differences was performed in SigmaStat 4.0 (Systat Software Inc., California, USA). Recordings of changes in the membrane potential were recorded with 2 Hz data collection frequency. Figures were prepared with the use of Sigma Plot 9.0 (Systat Software Inc., California, USA) and CorelDraw12 (Corel Corporation, Ottawa, Canada) software.

### Measurements of membrane changes in protonema cells

The plants were prepared in the same way as for the measurements carried out on the leaves. The microelectrode was positioned and inserted using a PatchStar (Scientifica, East Sussex, UK) micromanipulator and observed under an Olympus IX71 (Tokyo, Japan) microscope with a camera (Artcam-500MI, Tokyo, Japan) working with QuickPhoto Camera software (version 2.3, Promicra, Prague, Czech Republic). A CellTram Vario microinjector (Eppendorf, Hamburg, Germany) with a borosilicate glass micropipette (with a diameter of about 3 µm) was used for the application of H_2_O_2_ onto the cell surface. Likewise, in our previous paper (Koselski et al. 2020), 1 mM methyl blue was used for staining of the stimulating solution, which allows observation of its dispersion. Live recording of membrane potentials visible in LabScribe3 software and microscope camera images were recorded by OBS software (ver. 23.2.1; Open Broadcaster Software, Massachusetts, USA).

### Fluorescence calcium imaging

Films and images of calcium concentration changes in the GCaMP3 *P. patens* mutants were recorded with the use of NIS-Elements AR software (ver.5.20.00 Nikon, Tokyo, Japan) working with a Nikon Eclipse Ti fluorescence microscope equipped with a Nikon Plan UW 2X WD:7,5 objective and a Nikon DS-Ri2 camera (Nikon, Tokyo, Japan). The excitation of fluorescence was provided by a Prior lumen 200 metal arc lamp (Prior Scientific Instruments Ltd., Cambridge, England) and a 495-nm dichroic mirror with a standard GFP excitation filter 470 ± 20 nm. Images of fluorescence emission were recorded with 1-s exposure and a standard GFP fluorescence emission barrier filter 525 ± 50 nm (Nikon, Tokyo, Japan). Stimulation of the plants was carried out by microinjection using a CellTram Vario microinjector (Eppendorf, Hamburg, Germany). The standard solution supplemented with H_2_O_2_ was visible due to tinting by 0.025 µM fluorescein.

### Treatment of plants used for gene expression analysis

Gametophores from hydroponic cultures were used for the treatment with H_2_O_2_. Half of the gametophores from one glass ring were cut approximately in the middle and only the upper half of the gametophores (∼100 mg) was used as control sample. Cut gametophore apices were gently dried by dabbing with filter paper. The glass ring with the remaining uncut gametophores was transferred into a new Magenta^®^Vessel containing KnopME with 0.5 mM H_2_O_2_. After eight minutes, the upper half of the remaining uncut gametophores (treated sample) was harvested as described before. In total, gametophores from four independent hydroponic cultures (four biological replicates) were sampled.

### RNA extraction and quantitative real-time PCR (qPCR)

Extraction of RNA was done with the innuPREP Plant RNA Kit (Analytic Jena, Jena, Germany) using the extraction buffer “PL”. 5 µg total RNA were treated with DNAseI (Thermo Fisher Scientific, Waltham, Massachusetts, U.S.) at 37°C for 1 hour and integrity of the RNA was checked on Agarose gels. 2 µg DNAseI digested RNA were used for reverse transcription using the TaqMan^TM^ Reverse Transcription kit (N8080234, Thermo Fisher Scientific) with random hexamer primers. Reverse transcription was performed at 42°C for 1 hour. A non-transcribed control without the addition of MultiScribe^TM^RT enzyme was included. Primers for the qPCR were designed using Primer3Plus software (https://www.primer3plus.com/, Untergasser et al., 2012) with qPCR settings and an efficiency of 2 was confirmed using a 1:2 dilution series of cDNA. Melting curve analysis was performed to exclude the presence of off-targets. qPCR was performed in 96-well plates using the SensiFast^TM^ SYBR No-ROX Kit (Bioline) in technical triplicates for each biological replicate. 50 ng cDNA were used for each technical triplicate and the PCR reaction was performed in a LightCycler^®^ 480 (Roche, Basel, Switzerland). -RT and water controls for each primer pair were included. The PCR reaction was performed in 45 cycles with a melting temperature of 60°C. Expression analysis of the genes Pp3c5_28420V3 and Pp3c16_25090V3 (genes of interest, GOI) was performed as described (Bohlender et al., 2020) in relation to the housekeeping (reference) genes L21 (Pp3c13_2360V3, Beike et al., 2015) and LWD (Pp3c22_18860V3, Schuessele et al., 2016). All primers are listed in Supplementary Table 1. The expression levels were calculated relative to the reference genes according to Livak and Schmittgen (2001) using the software for the LightCycler® 480 (V1.5.0, Roche). Relative expression is represented as 2^(-ΔCT)^ with ΔCT = CT_[GOI]_ – CT_[reference]_. Figure 6 was created and statistics were calculated in R (R Core Team, 2022). Statistical significance was tested via one-way Anova with subsequent post-hoc test. Significance was accepted at p < 0.05.

## Data Availability

The data underlying this article are available in the article and in its online supplementary material.

## Acknowledgments

The authors thank Prof. José A. Feijó, Dr. Jörg D. Becker and Dr. Mário R. Santos for sharing the glutamate receptor 1 (GLR1) knockout mutant of Physcomitrella. The authors thank Dr. Thomas J. Kleist for providing the transgenic lines of Physcomitrella expressing GCaMP3. We thank Richard Haas for performing the qPCR experiments and Anne Katrin Prowse for language editing.

## Author contribution

M.K. conceived and designed the manuscript, prepared the main part of experiments, analyzed and interpreted the data, and drafted the main part of the work. P.W. participated in preparation of the membrane potential measurements. K.T. participated in writing and revised the manuscript. S. N. W. H and R. R. designed qPCR experiments, analyzed data and helped writing the manuscript.

