## Supplementary Figures for "Long-Distance Electrical and Calcium Signals Evoked by Hydrogen Peroxide in Physcomitrella"

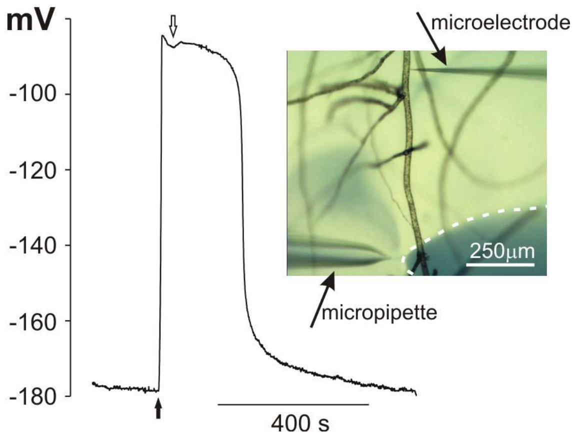


**Supplementary Figure 1.** Membrane potential changes recorded after application of 0.5 mM hydrogen peroxide to the protonema cells adjacent to the tested cell (site of the microelectrode insertion). The tested cell was located in the same chain of cells as stimulated cells. The picture placed on the right shows dispersion of $H_2O_2$ stained with 1 mM aniline blue (marked by dashed white lines) recorded 5 seconds after stimulation marked on the trace by a black arrow. The end of stimulation (removal of the micropipette from the measuring chamber) is marked by white arrow.

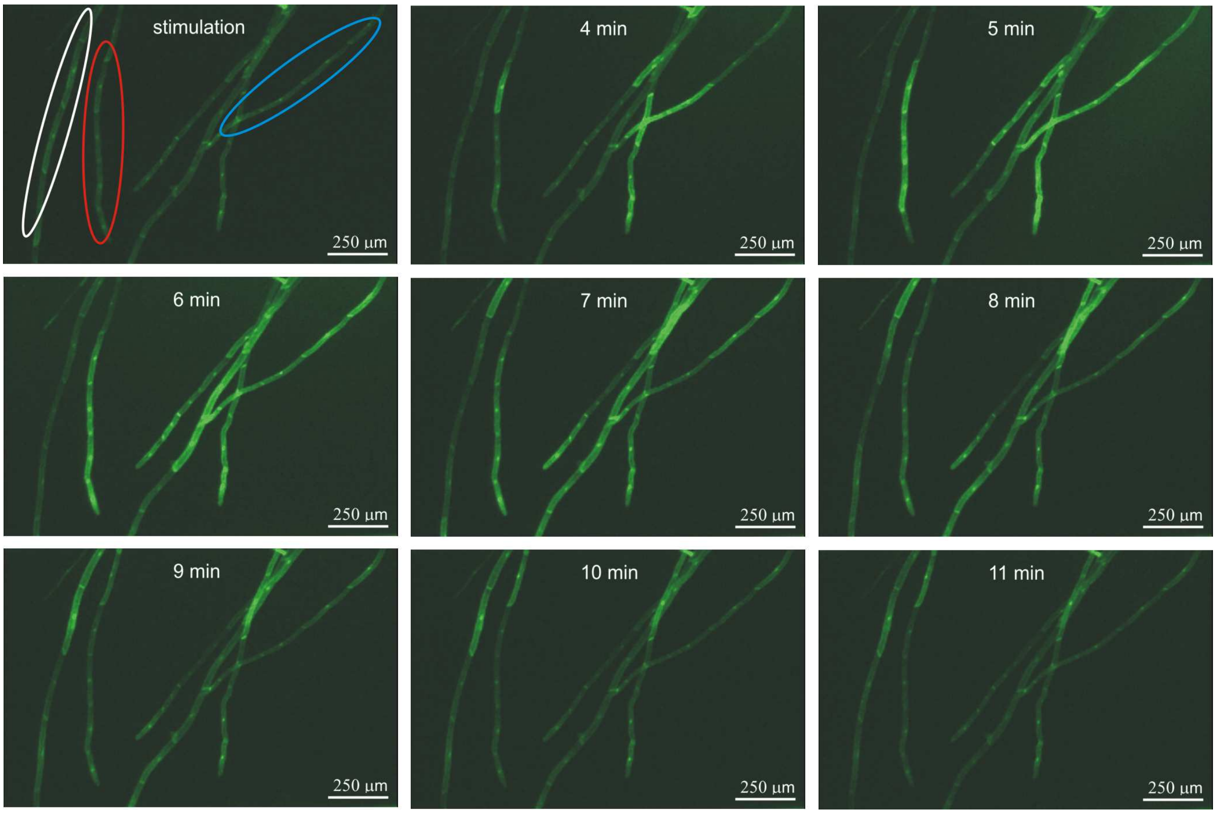


**Supplementary Figure 2.** Pictures presenting differences in calcium signal propagation recorded in three different chains of protonema cells. The experiment was carried out on the cells expressing fluorescent calcium biosensor GCaMP3. The responses were evoked by application of 0.5 mM hydrogen peroxide to the basal part of the gametophyte. Blue and red ellipse indicate two directions of calcium signal propagation - from the place of stimulation and in the opposite direction, respectively. White ellipse show the protonema cells in which decrement in calcium signal propagation was observed.

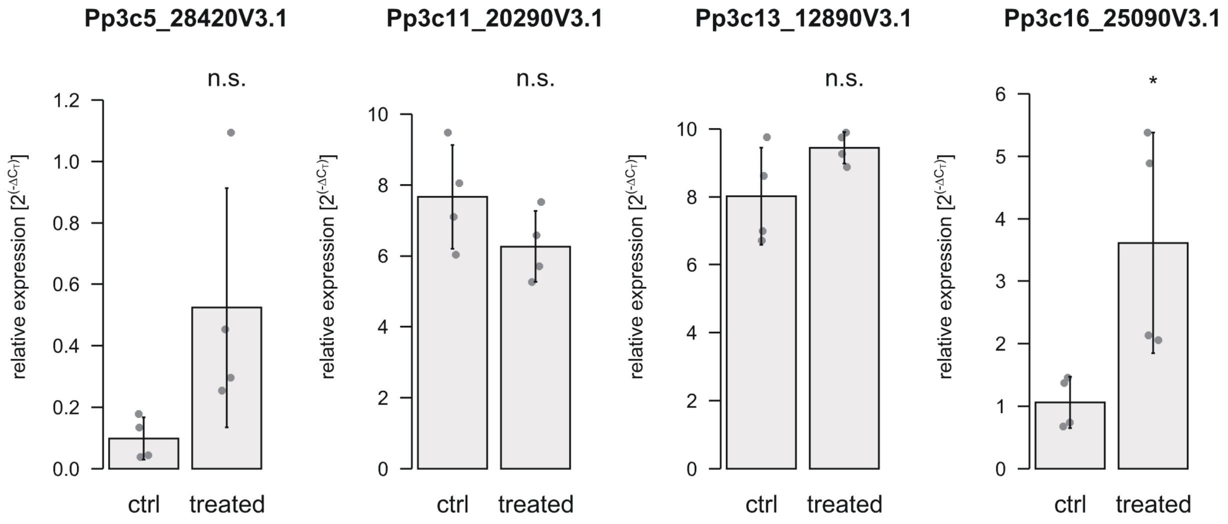


**Supplementary Figure 3.** Gene expression analysis by qPCR of four selected gene candidates (Pp3c5_28420V3.1, Pp3c25_14930V3.1, Pp3c16_25090V3.1, Pp3c2_18670V3.1). Gametophores from hydroponic cultures were submerged in 0.5 mM $H_2O_2$ for 8 min (treated). Dots represent biological replicates (mean values from three technical replicates). Bars are mean values from the four biological replicates with standard deviation. Relative expression ($2^{(-\Delta CT)}$) of the candidates is calculated against the reference genes L21 (Pp3c13_2360V3.1) and LWD (Pp3c22_18860V3.1) according to Livak & Schmittgen (2001). Significance levels are based on a one-way Anova with subsequent post-hoc test (*p < 0.05); n.s. = not significant.

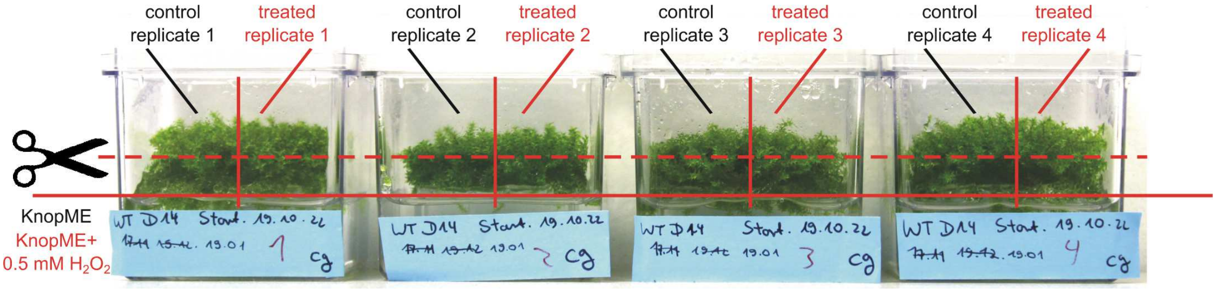


**Supplementary Figure 4.** Experimental setup for local treatment (base) of gametophores to analyze gene expression in the apex. Gametophores were cultivated on mesh fixed on glass rings. From four independent hydroponic gametophore cultures, the apices of approximately half of the gametophores were cut with scissors and kept as control samples (~100 mg fresh weight). The glass ring containing the remaining (uncut) gametophores was transferred to a new box containing KnopME with 0.5 mM $H_2O_2$ and incubated for 8 min. The level of the medium was always adjusted until it reached the bottom of the mesh (control and treatment). Apices of the remaining gametophores were harvested as treated samples.

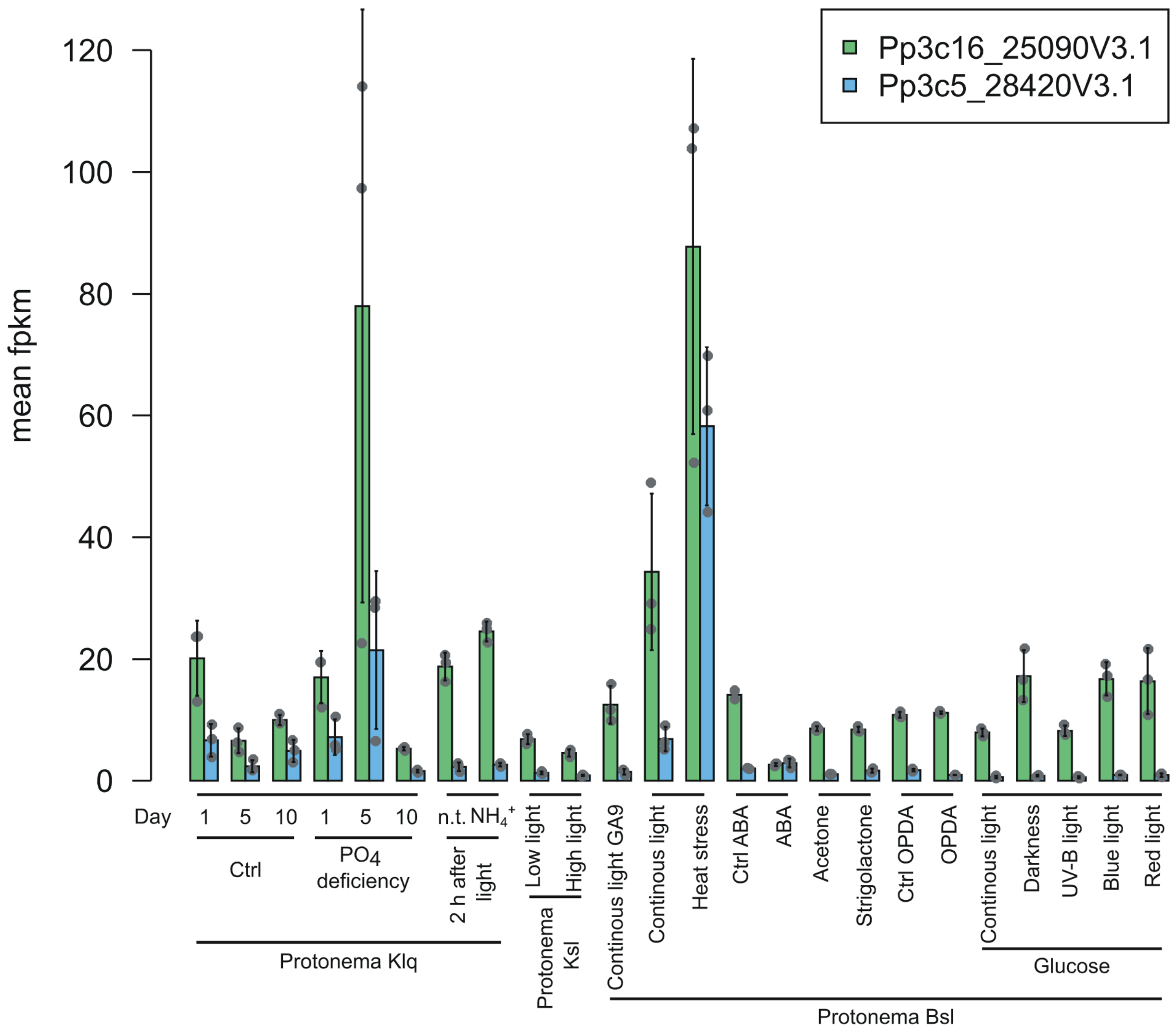


**Supplementary Figure 5**. Expression levels of the genes Pp3c5_28420V3.1 and Pp3c16_25090V3.1 after different treatments. All data were downloaded from PEATmoss (https://peatmoss.plantcode.cup.uni-freiburg.de/) and are described in Perroud et al. (2018) and Fernandez-Pozo et al. (2020). Mean FPKM (Fragments Per Kilobase Million) values with standard deviation are depicted. Data points represent biological replicates. In the case of the ABA control (Ctrl ABA) only data from two replicates was available. Abbreviations: B = BCD medium; lq = liquid; sl = solid; K = Knop medium; n.t. = not treated.
